# HIV-1 Rev-RRE Functional Activity in Primary Isolates is Highly Dependent on Minimal Context-Dependent Changes in Rev

**DOI:** 10.1101/2022.04.29.490105

**Authors:** Godfrey Dzhivhuho, Jordan Holsey, Ethan Honeycutt, Heather O’Farrell, David Rekosh, Marie-Louise Hammarskjold, Patrick E. H. Jackson

## Abstract

During HIV infection, intron-containing viral mRNAs have to be exported efficiently from the host cell nucleus to the cytoplasm in order to complete the replication cycle. To overcome cellular restrictions to export incompletely spliced transcripts, HIV encodes a protein, Rev, that is constitutively expressed from a completely spliced transcript. Rev is then imported into the nucleus where it binds to an RNA structure on intron-containing viral mRNAs called the Rev Response Element (RRE). Bound Rev multimerizes and recruits cellular factors that permit the nuclear export of the resulting ribonucleoprotein complex. Primary HIV isolates display substantial variation in the functional activity of the Rev-RRE axis, which may permit viral adaptation to differing immune environments. We describe two subtype G primary isolates with disparate Rev activity. Rev activity was correlated with *in vitro* fitness of replication-competent viral constructs. Amino acid differences within the oligomerziation domain, but not within the arginine-rich motif or nuclear export signal, determined the different levels of Rev activity. Two specific amino acid substitutions were demonstrated to be able to alter the low-activity Rev to a high-activity phenotype. However, introducing the original amino acids from the the low activity Rev into high activity Rev in this position did not result in significant alterations in activity, highlighting the importance of the broader sequence context for functional activity. These results demonstrate that studies of Rev and RRE activity variation, which may have broader implications for HIV transmission and pathogenesis, should include sequences from primary isolates, as findings using only laboratory-adapted strains cannot be generalized.

## Introduction

Retroviruses must export intron-containing viral mRNAs from the cell nucleus to the cytoplasm in order to complete their replication cycles. So-called “complex” retroviruses utilize a trans-acting protein and a *cis*-acting RNA secondary structure to overcome the cellular restriction on the export of unspliced and incompletely spliced transcripts (1). The complex retrovirus HIV-1 requires both the viral protein Rev and an RNA structure termed the Rev-Response Element (RRE) to accomplish this process (2, 3). During HIV replication, Rev is translated from a completely spliced, and thus constitutively exported, mRNA species (4–6). After translation, Rev is imported into the nucleus where it binds to the RRE found on all unspliced and incompletely spliced viral transcripts (7–10). A Rev homo-oligomer forms and then recruits cellular factors including Crm1 and Ran-GTP. The resulting ribonucleoprotein complex is then exported to the nucleus where the intron-containing viral mRNA can be translated or packaged into progeny viruses (11–13).

Primary isolates of HIV exhibit sequence variation throughout the genome, including in the regions encoding *rev* and the RRE (14). Variations in both *rev* and the RRE are observed not only between primary isolates sequenced from different hosts, but also within viral quasispecies arising during the course of infection in a single host. Sequence changes in *rev*, the RRE, or both of these elements can give rise to differences in the functional activity of the Rev-RRE regulatory axis (15–18).

The Rev gene in subtype B laboratory isolates that have been used in most studies to date encodes a 116 amino acid protein. Rev contains several functional domains, separated by less ordered spacer regions. The core functional domains are the bipartite oligomerization domain (OD), the arginine-rich motif (ARM), and the nuclear export signal (NES). The OD consists of two alpha-helical regions which stabilize the Rev monomer and permit the formation of homodimers and higher order Rev oligomers (19). The ARM is the site of direct interaction with the RRE and doubles as a nuclear localization signal (20–23). The NES permits Rev oligomer interaction with Crm1 to accomplish nuclear export (3, 24). The N-terminal and C-terminal regions of Rev are less well ordered. A “Turn” region is located between the first portion of the OD and the ARM, and a “Link” region is located between the second portion of the OD and the NES (25).

We previously described twelve HIV-1 primary isolates that displayed substantial variation in the activity of the Rev-RRE regulatory axes (15). Within this set, we found two subtype G viruses with widely disparate Rev-RRE activity, attributable to differences between the Rev proteins rather than the RREs. Furthermore, the difference in Rev activity was not due to a difference in steady-state protein level, suggesting that the different sequences resulted in proteins with intrinsically different levels of activity.

Previous work describing differential Rev activity in primary isolates has been restricted to studies of subtype B and C viruses. Rev and RRE variation in other HIV subtypes has not been well characterized. In this study, we sought to identify the key sequence determinants of Rev functional activity, using the two subtype G primary isolates as a model.

## Results

### Determination of native Rev functional activity

We previously described the relative functional activity of the two subtype G Revs using a transient transfection-based lentiviral vector assay system (described in (18)). In that system, vector RNA was packaged in a Rev-RRE dependent fashion and the resulting vector titer corresponded to the relative activity level of the Rev-RRE pair Using this assay, the 8-G Rev/NL4-3 RRE pair was found to have notably low functional activity, while the 9-G Rev/NL4-3 RRE pair had high functional activity (15). The finding of differential Rev activity was consistent when the two Revs were paired with the RRE from either the 8-G or 9-G primary isolate as well.

To validate these results, a second assay system for determining Rev-RRE functional activity was used that better reproduces the natural process of viral infection. This fluorescence-based system permits the measurement of activity from integrated proviral chromatin in lymphoid cells, as opposed to using transiently transfected plasmid DNA. This assay was described fully in (26). Using this system, the relative functional activity of 8-G Rev was again found to be significantly lower than that of the 9-G Rev in combination with the NL4-3 RRE (*p* < 0.001) (Figure 1A). This difference in activity persisted when the Revs were tested in combination with the RREs from the 8-G and 9-G primary isolates (*p* < 0.001 for both comparisons).

**Figure 1.**
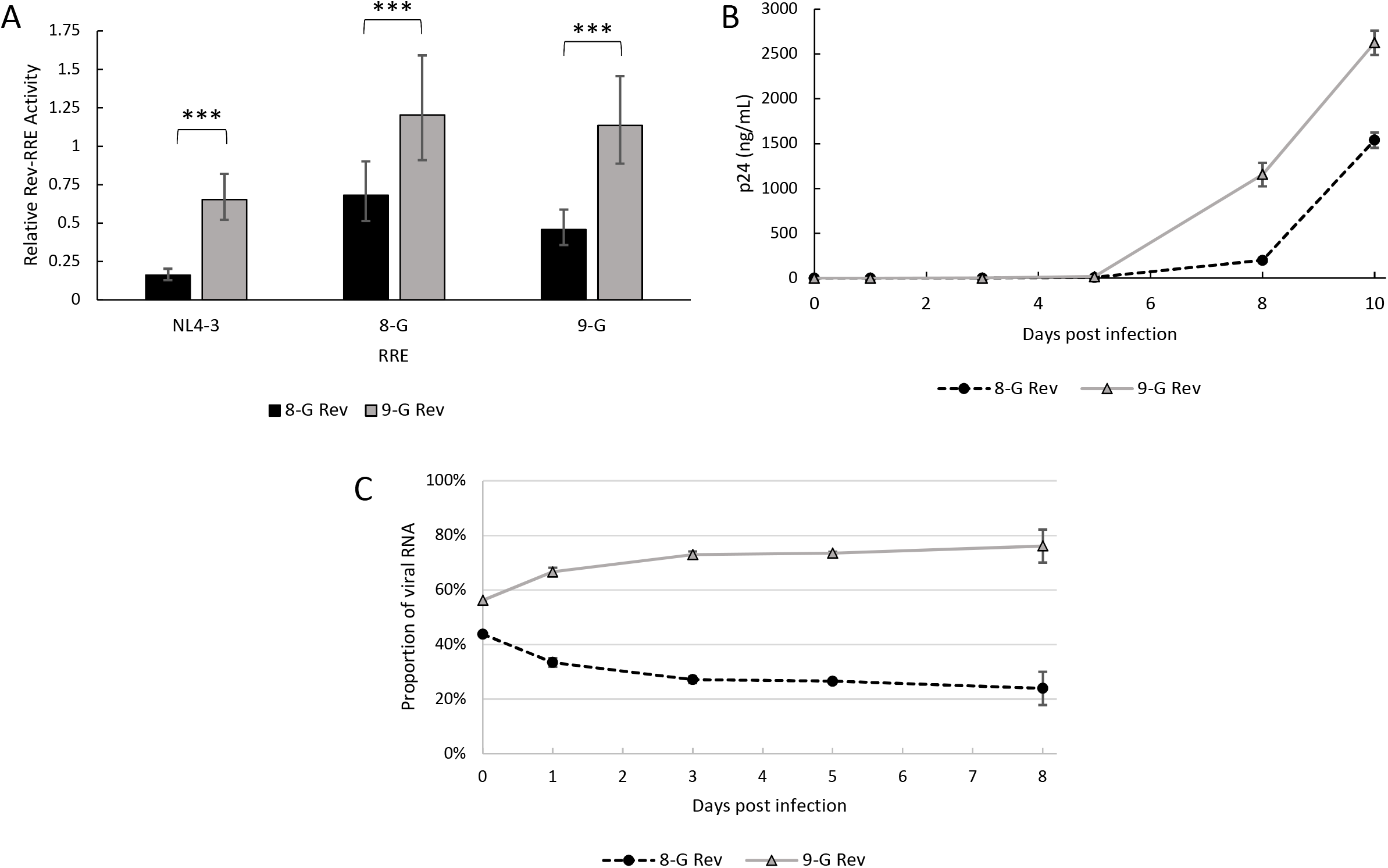
Relative activity of the 8-G and 9-G Revs. The activity of the Revs derived from the 8-G and 9-G primary isolates was determined *via* three complementary methods. A. Rev-RRE activity was determined by a fluorescence-based assay utilizing lymphoid cells transduced with Rev- and RRE-containing reporter constructs. The relative activity of the 8-G and 9-G Revs was measured in conjunction with the NL4-3, 8-G, and 9-G RREs. Relative activity is expressed in arbitrary units. *N* > 3 for all comparisons, error bars represent 95% confidence intervals (CI), *** *p* < 0.001. B. Replication-competent constructs containing the 8-G or 9-G Rev were used to infect parallel cultures of SupT1 cells. Infection kinetics were measured by determining the production of p24 protein in culture supernatants over time. *N* = 3 for each data point, error bars represent standard deviation (SD). C. The relative replicative fitness of the replication-competent constructs assayed in (B) was also measured in a direct competition assay. Cultures of SupT1 cells were co-infected with 8-G and 9-G Rev containing viruses. The relative amount of 8-G Rev and 9-G Rev sequence in viral cDNA was determined at multiple time points, and expressed as the percentage of total Rev cDNA at that time point. *N* = 3 for each data point, error bars represent SD.

To further confirm this finding, the activity of the two Revs was also tested in a spreading infection assay. The laboratory HIV strain NL4-3 was modified with two stop codons in the first coding exon of Rev (without altering the Tat sequence) to silence native *rev* expression, and a *rev-nef* cassette was inserted in the *nef* position with the two genes separated by an internal ribosomal entry site (IRES). This resulted in a replication-competent construct expressing all viral genes with an exchangeable *rev* in a heterotopic position. Viruses were created containing the 8-G or the 9-G *rev* and used to infect parallel cultures of SupT1 cells. Viral growth kinetics were determined by serial measurements of p24 starting on the day of infection. The virus containing 9-G rev yielded a substantially higher amount of p24 at an earlier time after infection than did the virus containing 8-G *rev*, consistent with greater functional activity (Figure 1B).

Finally, the difference in viral replication kinetics due to the different *rev* sequences was verified by a direct competition assay. Cultures of SupT1 cells were co-infected with the 8-G and 9-G rev-containing viruses. After infection, a portion of the culture supernatant was collected at serial time points and viral mRNA was prepared. The relative amount of 8-G and 9-G viral RNA at each time point was determined by PCR. The ratio of 9-G construct RNA to 8-G construct RNA increased over time, consistent with greater fitness of the 9-G *rev* containing virus (Figure 1C).

### Creation of chimeric Rev constructs

The 8-G and 9-G Rev sequences differ from each other at 29 amino acid positions, with substitutions in all functional regions and a seven amino acid deletion in the 9-G c-terminal region relative to 8-G (Figure 2). Chimeric *rev* sequences were designed for use in the fluorescence-based functional assay consisting of either the 8-G or 9-G sequence with one or more functional domains swapped for the sequence corresponding to the other Rev. For example, the “8-G with 9-G Turn” rev was identical to 8-G except for the Turn region (aa 26 to 34), the sequence of which was identical to the corresponding region from 9-G. Additional sequences were designed with single or multiple amino acid substitutions at key positions. Designed rev sequences were cloned into packageable constructs for the performance of functional assays.

**Figure 2.**
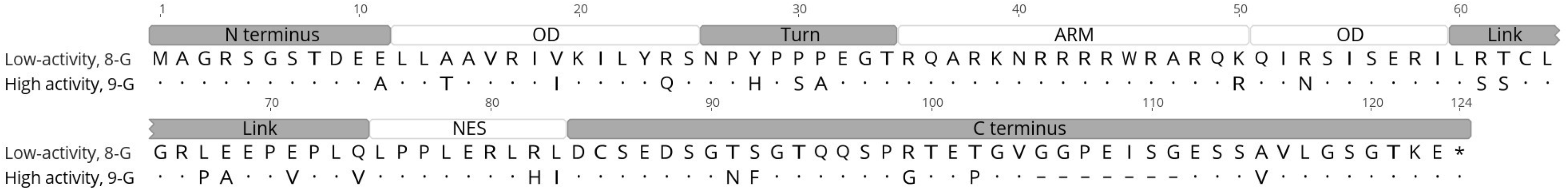
Alignment of 8-G and 9-G Revs. The amino acid sequences of the 8-G and 9-G Revs were aligned. Differences in the 9-G sequence relative to 8-G are highlighted. In the 9-G sequence, dots represent identical amino acid residues to the 8-G sequence, while dashes represent a deletion in the 9-G sequence relative to 8-G. Functional domains are marked above the alignment. OD – oligomerization domain, ARM – arginine rich motif, NES – nuclear export signal.

### Identification of Rev functional domains responsible for differential activity

To define the regions of Rev determining the difference in activity, chimeric sequences based on the 8-G rev were first tested with the NL4-3 RRE in the functional activity assay (Figure 3A). Two 8-G-based rev chimeras displayed increased activity relative to the native 8-G sequence: one with 9-G rev sequence in the N-terminus and the first portion of the OD (8-G with 9-G N+OD) and a second with 9-G *rev* sequence in the ARM and second portion of the OD (8-G with 9-G ARM+OD) (*p* < 0.001 for both comparisons). The 8-G *rev* chimeras with substituted Turn and NES regions or with a substituted Link region did not significantly differ in activity from the native 8-G *rev*.

**Figure 3.**
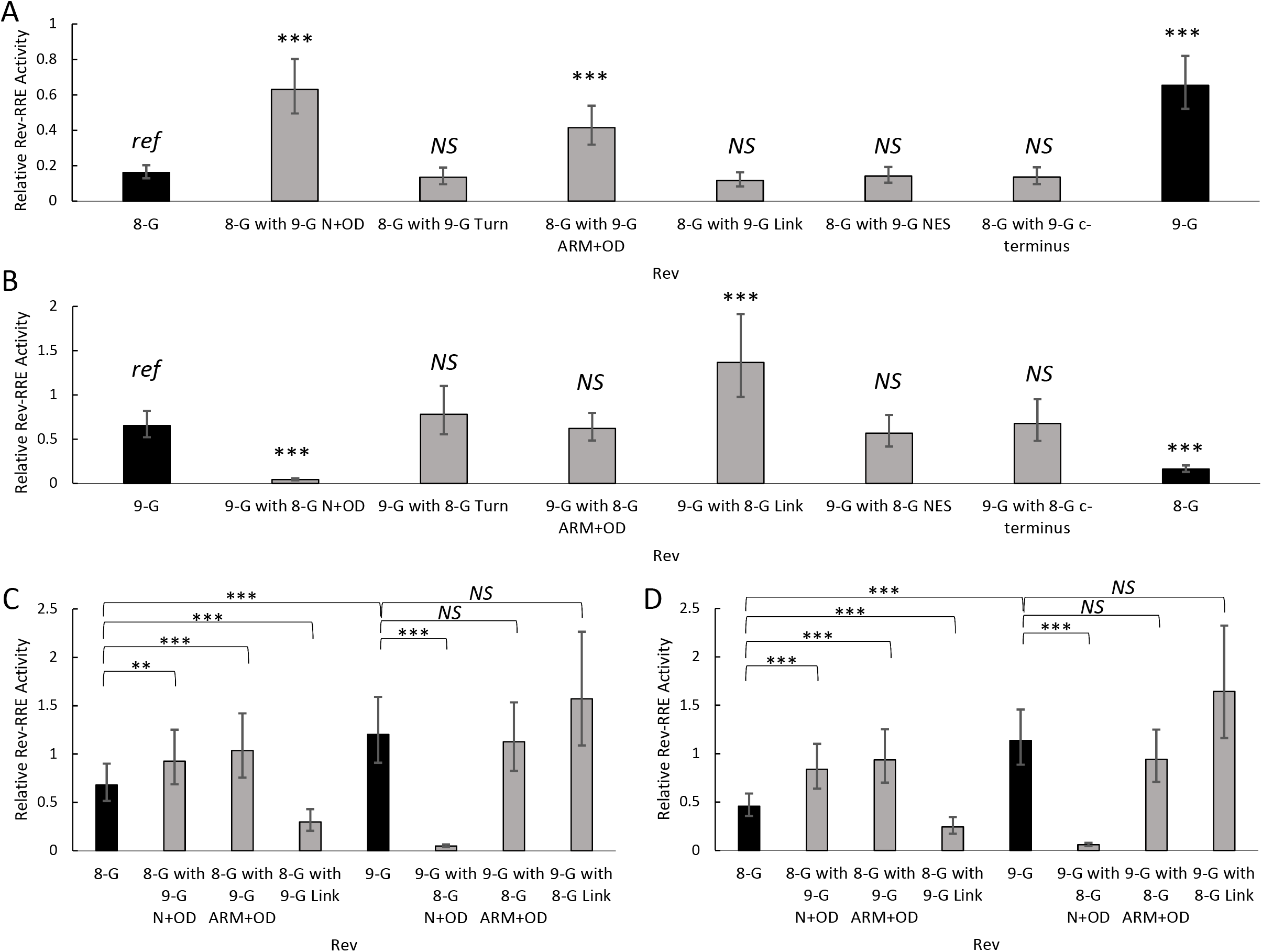
Activity of chimeric Rev sequences. Chimeras of 8-G and 9-G Rev were created by exchanging portions of the sequence corresponding to specific functional domains. Functional activity was determined using the fluorescence-based assay system. In each plot, the native 8-G and 9-G Rev sequences are included in dark bars for reference and chimeras are in light bars. A. 8-G chimeras tested on the NL4-3 RRE. B. 9-G chimeras tested on the NL4-3 RRE. C. Selected chimeras tested on the 8-G RRE. D. Selected chimeras tested on the 9-G RRE. For plots A and B, the statistical comparison is performed in reference to the left-most Rev, while in plots C and D the two Revs being compared are indicated by the horizontal bar. *N* ≥ 3 for all data points, error bars represent 95% CI. N+OD – n-terminus plus first portion of the oligomerization domain, ARM+OD – arginine rich motif plus second portion of the oligomerization domain. *Ref* – reference sequence for the plot, *NS* – not significant, *** *p* < 0.001, ** *p* < 0.01.

Complementary chimeras based on the 9-G *rev* sequence were also tested with the NL4-3 RRE (Figure 3B). Replacement of the 9-G n-terminus and first portion of the OD region with sequence from 8-G (9-G with 8-G N+OD) led to significantly lower functional activity than native 9-G *rev* sequence (*p* < 0.001), while replacement of the Link region resulted in greater functional activity (*p* < 0.001). In contrast with the 8-G-based chimera, substitution of the ARM and second portion of the OD did not alter functional activity in the 9-G context (*p* = 0.89).

Functional activity of the 8-G and 9-G *rev* chimeras was also tested with the 8-G RRE (Figure 3C) and 9-G RRE (Figure 3D). On these RREs, both the 8-G with 9-G N+OD *rev* and the 8-G with 9-G ARM+OD *rev* chimeras consistently displayed higher functional activity than native 8-G Rev (*p* < 0.001 for all comparisons). For the remaining *rev* chimeras, the pattern of activity alteration was similar across all three RREs.

Exchange of the c-terminal region did not significantly affect functional activity of either the 8-G or 9-G *rev* sequence on any of the three RREs (Figure 3). Replicating the seven amino acid insertion/deletion within the c-terminus also had no effect on activity, while truncation of the c-terminus after position 83 in both Revs significantly decreased functional activity relative to the corresponding native sequence (*p* < 0.001) (Figure S1).

### Functional activity alteration as assessed by single amino acid substitutions

Having identified the N-terminal, OD, and ARM domains as key for defining Rev functional activity, mutants with single amino acid changes within these regions were next created and tested on the NL4-3 RRE (Figure 4).

**Figure 4.**
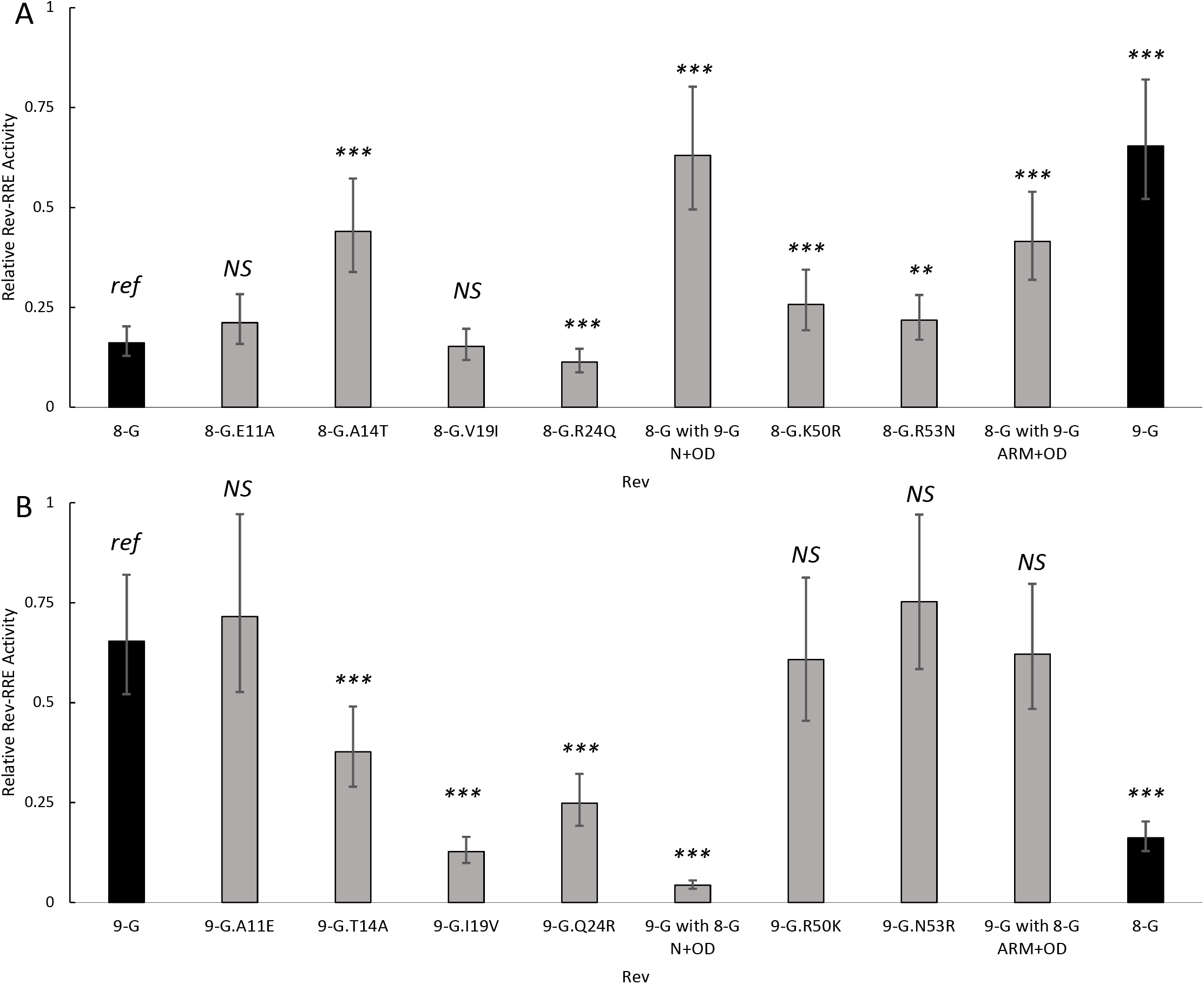
Activity of Revs with single amino acid substitutions on NL4-3 RRE. 8-G (A) and 9-G (B) Revs with single amino acid substitutions within the N-terminal, OD, or ARM regions were created and their functional activity determined on the NL4-3 RRE using the fluorescence-based assay system. In each plot, the native 8-G and 9-G Rev sequences are included in dark bars for reference and modified Revs are in light bars. The statistical comparison is performed in reference to the left-most Rev in each plot. *N* ≥ 3 for all data points, error bars represent 95% CI. *Ref* – reference sequence for the plot, *NS* – not significant, *** *p* < 0.001, ** *p* < 0.01.

Four amino acid differences between the native 8-G and 9-G Rev sequences occurred in the N-terminus and first portion of the OD, at positions 11, 14, 19, and 24. Only the 8-G.A14T mutant displayed significantly greater activity than the native 8-G Rev (*p* < 0.001). The 8-G.E11A and 8-G.V19I mutants displayed similar activity to 8-G Rev, while the 8-G.R24Q mutant displayed significantly lower activity (*p* = 0.001). Reciprocal substitutions in the 9-G Rev context showed a significant decrease in activity for the 9-G.T14A, 9-G.I19V, and 9-G.Q24R mutants relative to native 9-G Rev, while the 9-G.A11E substitution did not change activity.

Two amino acid differences occurred within the ARM and second portion of the OD at positions 50 and 53. Both the 8-G.K50R and the 8-G.R53N substitutions increased functional activity over native 8-G Rev (*p* < 0.01). The corresponding substitutions 9-G.R50K and 9-G.N53R did not alter activity relative to native 9-G.

A similar pattern in activity level was seen when the Rev mutants were tested on the 8-G and 9-G RREs (Figure S2). The 8-G.A14T Rev displayed consistently greater activity than the native sequence, while 8-G.R24Q displayed consistently lower activity (*p* ≤ 0.01 for all comparisons).

### Functional activity alteration with multiple amino acid substitutions

Rev mutants incorporating combinations of amino acid substitutions were created to identify the minimum change required to define the activity phenotype (Figure 5). While the 8-G.A14T substitution was sufficient to significantly increase the activity of 8-G Rev over the native sequence, the mutant was still less active than the native 9-G Rev (*p* < 0.001). The 8-G mutants 8-G.A14T+V19I and 8-G.A14T+R24Q, however, displayed functional activity that was similar to 9-G on the NL4-3 RRE. This pattern of activity was replicated with the 8-G and 9-G RREs (Figure S3).

**Figure 5.**
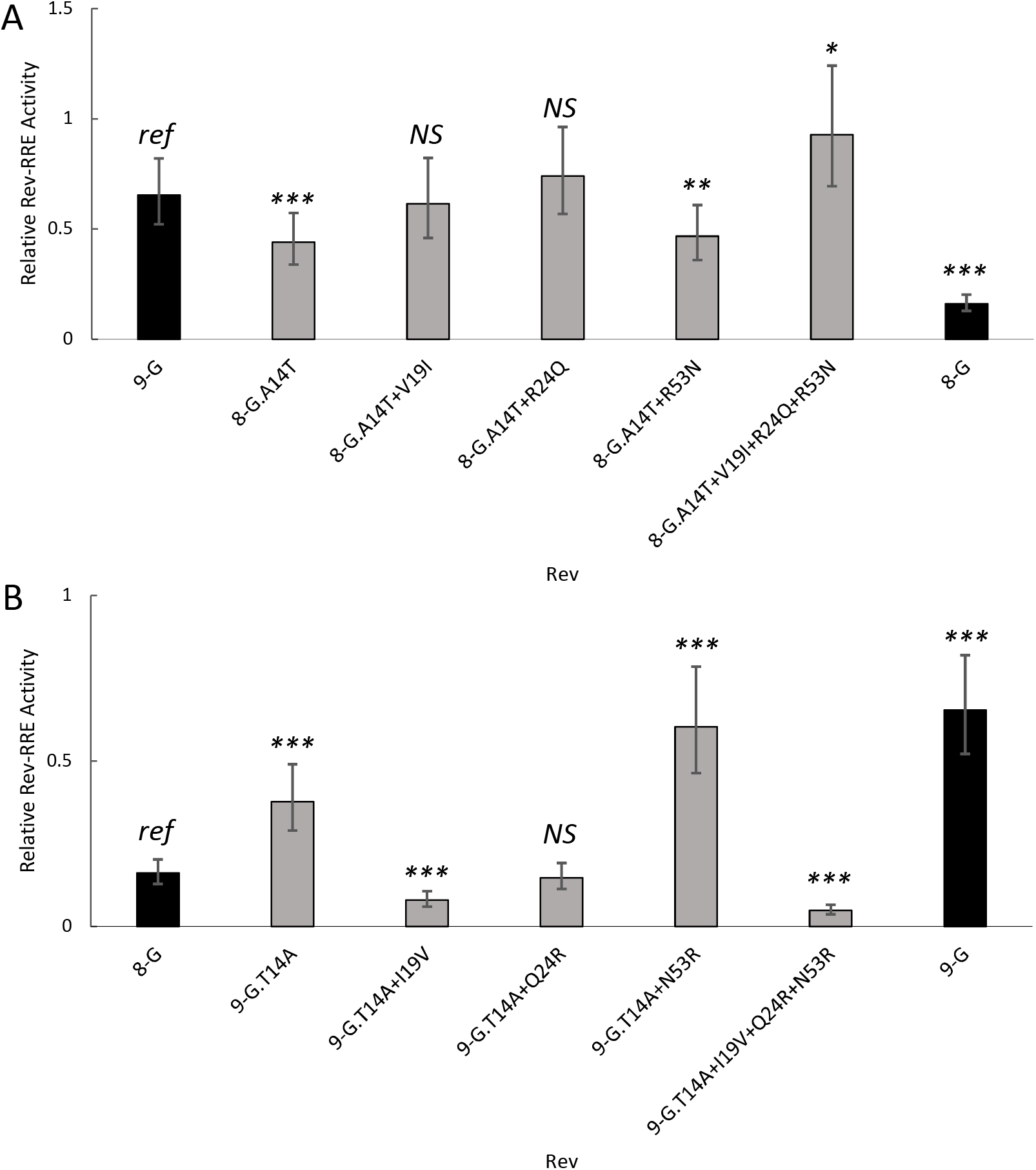
Activity of Revs with multiple amino acid substitutions on NL4-3 RRE. 8-G (A) and 9-G (B) Revs with multiple amino acid substitutions were created and their functional activity determined on the NL4-3 RRE using the fluorescence-based assay system. In each plot, the native 8-G and 9-G Rev sequences are included in dark bars for reference and modified Revs are in light bars. The statistical comparison is performed in reference to the left-most Rev in each plot. *N* ≥ 3 for all data points, error bars represent 95% CI. *Ref* – reference sequence for the plot, *NS* – not significant, *** *p* < 0.001, \*\**p* < 0.01, * *p* <0.05.

9-G Rev mutants containing the corresponding amino acid substitutions were also created and tested. On the NL4-3 RRE, the mutants 9-G.T14A+Q24R and 9-G.T14A+I19V displayed similar or lower activity than the native 8-G Rev. This pattern held for the 8-G and 9-G RREs as well.

### Effect of phosphomimetic substitutions

The amino acid residue at position 14 was found to be a key determinant of activity. One potential explanation for the increase in 8-G activity with the A14T substitution (and corresponding decrease in 9-G activity with the T14A substitution) could be linked to the potential for threonine, but not alanine, to become phosphorylated. The amino terminus of Rev is phosphorylated *in vivo* (27, 28), and Rev phosphorylation at other sites (but not position 14) has been associated with differences in binding to the RRE (29). To test whether phosphorylation of 14T could affect activity, phosphomimetic substitution mutants were created at this position (30). A 14S substitution presents an alternative phosphorylation target, while 14D and 14E substitutions structurally mimic phosphothreonine. The 14A substitution constitutes a site without potential for phosphorylation.

On the NL4-3 RRE, the phosphomimetic 8-G.A14S, 8-G.A14D, and 8-G.A14E mutants displayed similar or lower activity to the native 8-G sequence, and only the 8-G.A14T mutant showed higher activity (Figure 6). In the 9-G context, the 9-G.T14S, 9-G.T14D, and 9-G.T14E mutations did not significantly change 9-G activity and only 9-GT14A mutant activity differed from the native sequence. A similar pattern of activity was seen on the 8-G and 9-G RREs (Figure S4). Thus, while the phosphomimetic substitutions in the 9-G Rev context preserved a high level of activity, none of the corresponding substitutions in the 8-G Rev context increased activity similarly to that displayed by 8-G.A14T. Phosphorylation of position 14 is thus unlikely to explain the observed differences in functional activity.

**Figure 6.**
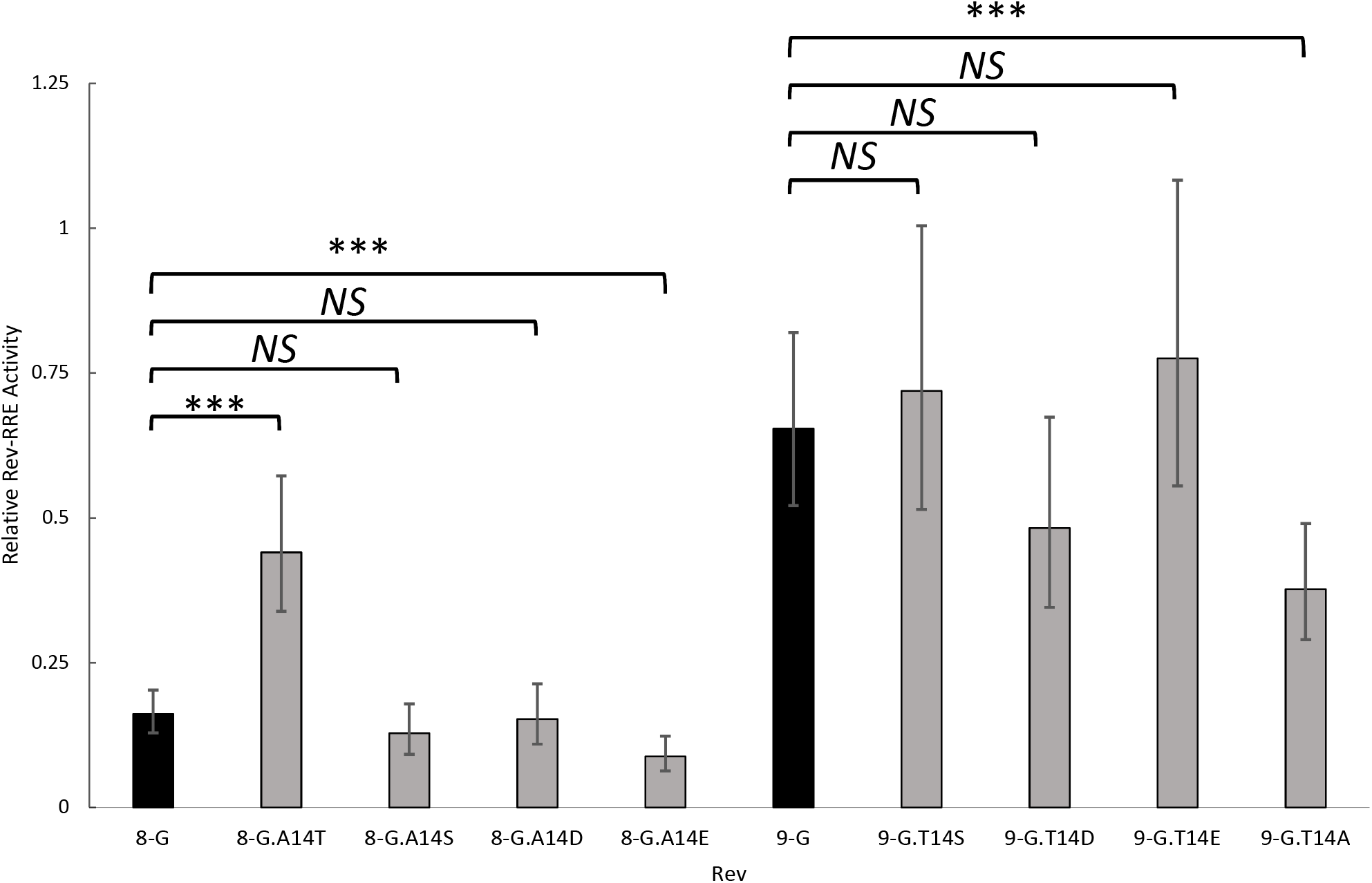
Activity of Revs with phosophomimetic substitutions on NL4-3 RRE. Revs with phosphomimetic amino acid substitutions at position 14 were created and their functional activity determined on the NL4-3 RRE using the fluorescence-based assay system. The native 8-G and 9-G Rev sequences are included in dark bars for reference and modified Revs are in light bars. The statistical comparison is performed is performed between the Revs indicated by the horizontal bars. *N* ≥ 3 for all data points, error bars represent 95% CI. *NS* – not significant, *** *p* < 0.001.

### Structural predictions of 8-G and 9-G Rev

Predicted structures of the native 8-G and 9-G Revs were computed using AlphaFold Colab (31) and visualized using PyMOL Molecular Graphics System, version 2.5.1 (Schrödinger, LLC). The resulting predictions were compared to the NL4-3 Rev dimer crystal structures previously published by Frankel and coworkers (32). The predicted subtype G structures overlapped each other and the NL4-3 crystal structure. Isoleucine 19, conserved in the NL4-3 Rev and 9-G Rev sequences, is positioned between the two OD helices within the Rev monomer, and in the RNA-bound crystal structure is expected to interact with I53, I55, and S56 residues found in all three Revs. In the 8-G Rev sequence, position 19 is a valine which may not be able to support this monomer-stabilizing interaction. We hypothesized that this difference could account for the decreased activity of the 9-G.I19V mutant relative to native 9-G as well as the increased activity seen with the 8-G.A14T+V19I mutant relative to 8-G.A14T.

To test this hypothesis, 8-G and 9-G mutants with a 19L substitution were generated and tested on the NL4-3 RRE (Figure 7). While the 8-G.V19I substitution alone did not alter activity relative to native 8-G, the 8-G.V19L mutant displayed significantly greater activity (*p* < 0.001). In the 9-G context, both 9-G.I19V and 9-G.I19L displayed lower activity than the native 9-G sequence (*p* < 0.001 for both comparisons), but 9-G.I19L displayed greater activity than 9-G.I19V (*p* < 0.001). A similar pattern of activity was seen with the 8-G and 9-G RREs (Figure S5). Thus, for both Revs the 19L variants display consistently greater activity than the 19V variants. However, the 19I variant yields low activity in the 8-G context but high activity in the 9-G context. The consequence of single amino acid substitutions in Rev at this position appears highly dependent on sequence differences elsewhere in the protein.

**Figure 7.**
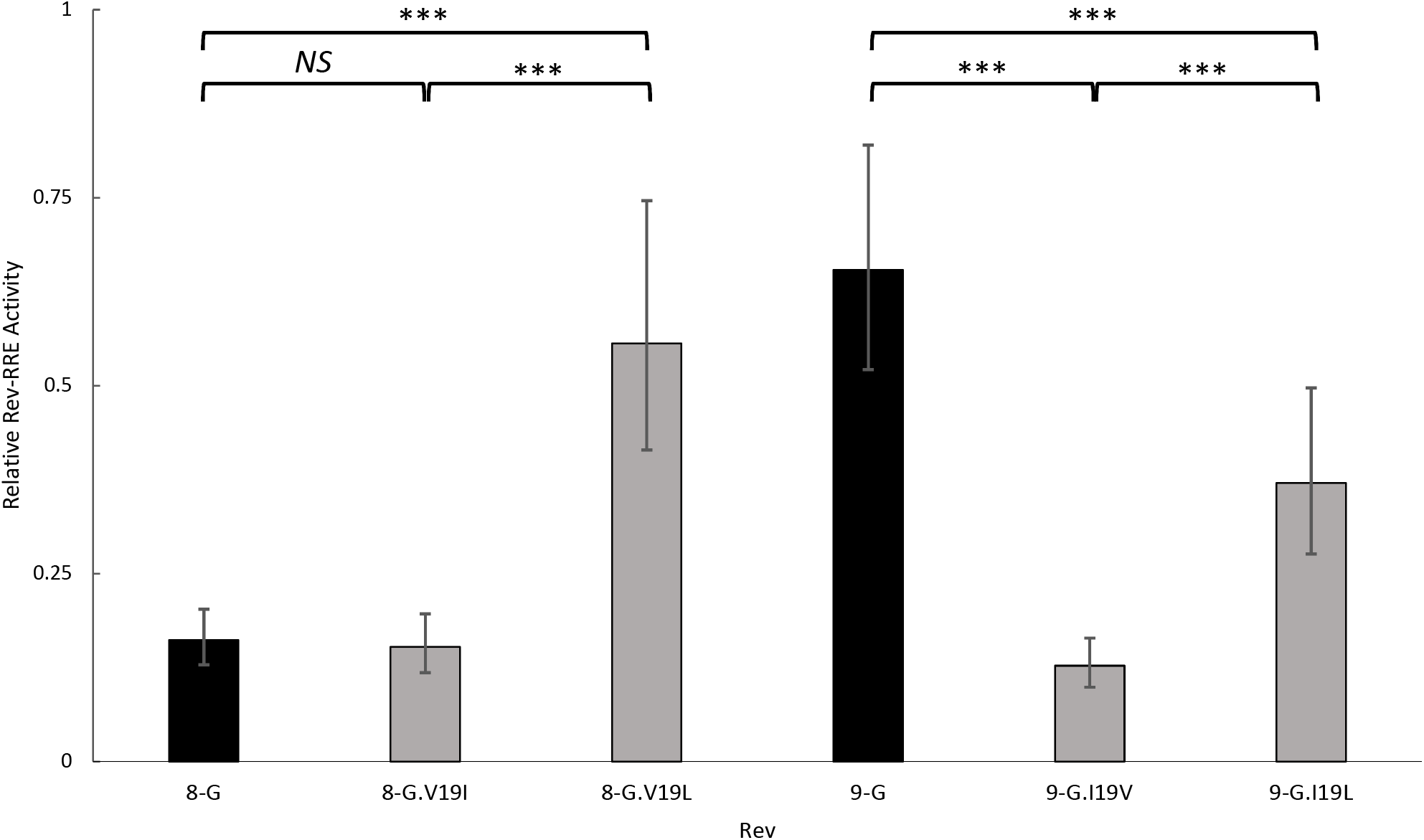
Activity of position 19 mutants on NL4-3 RRE. Revs with amino acid substitutions at position 19 were created and their functional activity determined on the NL4-3 RRE using the fluorescence-based assay system. The native 8-G and 9-G Rev sequences are included in dark bars for reference and modified Revs are in light bars. The statistical comparison is performed is performed between the Revs indicated by the horizontal bars. *N* ≥ 3 for all data points, error bars represent 95% CI. *NS* – not significant, *** *p* < 0.001.

## Discussion

This report adds to the sparse literature on the functional implications of naturally occurring variation in the HIV-1 Rev-RRE regulatory axis. To our knowledge, this is the first functional exploration of Rev variation in subtype G primary isolates. Using two Rev sequences with markedly different levels of activity, we demonstrated that amino acid differences within the oligomerization domain alone result in significant variation in Rev activity. Furthermore, just two amino acid substitutions in this region are sufficient to alter a low-activity to a high-activity Rev phenotype.

Modulation of the activity of the Rev-RRE axis may be a mechanism by which HIV adapts to an array of selection pressures that the virus encounters throughout the course of natural infection. The virus may replicate in the environment of the genital or rectal mucosa during transmission, in various anatomical compartments including blood and lymphoid tissue during chronic infection, and in conditions of attenuated immune surveillance during late disease. Experimental evidence suggests that low Rev-RRE functional activity is associated with evasion of T-cell mediated killing (33), while high RRE activity may result in a more rapid decline in CD4 count (34, 35). The ability to modulate replication kinetics and the presentation of viral antigens through differences in Rev-RRE functional activity may offer an advantage to viruses in differing fitness landscapes.

Previous reports of Rev variation in primary isolates are limited to subtype B (36) and C (37) examples. In these subtypes, changes in the Rev oligomerization domain (38, 39), NES (40, 41), and c-terminal region (42) have been associated with differences in activity. In a subtype B context, mutations at amino acid position 18 within the first OD were found to significantly alter Rev-dependent reporter gene expression independent of changes in oligomerization on the RRE (38). Our results in subtype G isolates are consistent with the finding that the Rev OD may be an important site for activity modulation.

The specific amino acid substitutions that account for activity level in these subtype G sequences, at positions 14 and 19, have not been previously described. Fernandes and coworkers performed an analysis of Rev functional activity using the subtype B NL4-3 virus as a model (43). Each amino acid position in Rev was mutated, and the relative replicative fitness of viruses with Rev mutations was determined. In this model, an R24Q substitution resulted in a substantial increase in fitness; E11A and A14T resulted in a slight increase in fitness; V19I and R53N had no substantial change; and K50R showed a substantial decrease in fitness. These single amino acid substitutions had very different effects in the context of the 8-G sequence. For example, while our assay system replicates the finding from Fernandes that glutamine rather than arginine at position 24 yields significantly higher NL4-3 Rev activity (data not shown), in 8-G, R24Q displayed significantly lower activity than the corresponding native Rev. On the other hand, the data with Q24R in 9-G was consistent with the data of Fernandes and coworkers

Because the functional impact of individual amino acid substitutions is highly context-dependent, making predictions of function based on structure is challenging. The NL4-3 Rev crystal structure suggests that the residue at position 19 is functionally important due to a role in stabilizing the interaction of the two OD regions within the Rev monomer. While this hypothesis seems to hold true for the 9-G Rev, in which both isoleucine and leucine at position 19 yield higher activity than valine, it does not help to predict activity in the 8-G context, where at position 19 isoleucine and valine give similar low levels of activity. Thus, while structural studies of Rev utilizing NL4-3 or other laboratory strains of HIV can yield valuable insights, the functional significance of individual amino acid substitutions cannot be generalized between even closely related viruses. Mechanistic investigations of HIV must take a broader view of viral diversity and test hypotheses in primary isolates as well as laboratory strains.

A corollary of the highly context-dependent consequence of single amino acid substitutions is that combinations of substitutions have unpredictable implications for Rev functional activity. The effect of multiple substitutions is not merely additive. While 8-G.V19I had similar activity to native 8-G, the addition of substitution V19I to A14T increased 8-G activity more than the A14T substitution alone.

Our results suggest that Rev functional activity is highly sensitive to small sequence changes. We previously described HIV-1 primary isolates with variable Rev-RRE functional activity attributable to differences in Rev, the RRE, or both (15). Single nucleotide substitutions within the RRE can give rise to different two-dimensional structures that, in turn, have significant differences in functional activity (17). As the RRE is located within a region of the HIV genome that codes only for *env*, we considered that there may be a lower barrier for modulation of Rev-RRE functional activity through changes in the RRE, as this can be accomplished with a minimal number of silent mutations in *env*. On the other hand, *rev* overlaps with *tat, env*, or both over its entire length, which might be expected to constrain *rev* mutations that could lead to differences in Rev-RRE activity. Our study now demonstrates that, in HIV-1 primary isolates, one or two amino acid substitutions within the oligomerization domain of Rev is sufficient to cause substantial alteration in activity. Given this substantial flexibility to alter Rev activity with minimal sequence changes, we would thus expect to observe variations in both *rev* and the RRE that have the capacity to modulate Rev-RRE activity, without a preference for RRE mutations. The Frankel group has observed that the potential for *rev* sequence variation is further enhanced by the fact that the functional regions of *tat* and *rev* are spacially segregated, such that mutations within key domains of rev would not be expected to have significant functional implications *for tat*, and *vice versa* (43).

In this study, we limited our attention to correlations between Rev sequence and functional activity. While we can exclude some explanations for differences in Rev activity, specifically phosphorylation at position 14 and stabilization of the Rev monomer at position 19, the mechanisms behind the observed variations in activity have not been defined. While we previously found that the steady state protein level of 8-G exceeded that of 9-G (15), it is still formally possible that differences in protein turnover kinetics rather than intrinsic differences in Rev activity could account for the functional activity variations of mutants observed in these assays. From the standpoint of viral adaptation to different selection pressures, however, it is immaterial whether differential Rev-RRE activity is achieved via any of an array of potential mechanisms. A Rev mutation that accomplishes a decrease in Rev-RRE activity through decreased Rev protein level may be equally useful to the virus as a mutation that decreases the affinity of the Rev-RRE nucleoprotein complex for Crm1. While elucidating molecular mechanisms of Rev-RRE activity variation remains an important question, it is separate from the question of the relationship between *rev* sequence variation and the activity of the Rev-RRE axis.

In this study, we describe novel amino acid substitutions within the oligomerization domain of Rev that give rise to significant differences in Rev-RRE functional activity, specifically within the subtype G context. One can hypothesize that modulation of the Rev-RRE regulatory axis is one mechanism by which HIV can alter its replication kinetics and the relative expression of immune-activating and immune-modulatory viral proteins (33). This would give the virus a means to adapt to the differing immune milieus that it might encounter during the course of disease progression. Additional studies of this regulatory axis in primary isolates from different clinical senarios are needed to further define the potential role in the pathogenesis of HIV disease.

## Methods

### Rev and RRE sequence selection

The basic *rev* and RRE sequences for these studies were obtained from one of two primary isolates (8-G, Genbank: FJ389367 (44); 9-G, Genbank: JX140676 (45)) or from the laboratory strain NL4-3 (Genbank: U26942 (46)). The sequence corresponding to the *rev* or RRE from each viral genome was analyzed and extracted using Geneious Prime v11.0.9 (Biomatters).

### Replication competent constructs and viral stocks

Replication competent HIV constructs containing the 8-G and 9-G *rev* sequences were constructed to determine differences in replication kinetics. Full-length NL4-3 HIV (47) was modified to silence native *rev* expression by mutating the initial AUG codon to ACG and changing the codon at position 23 (UAU: Y) to a stop codon (UAA) without altering the sequence of Tat. The native *nef* was replaced, beginning at the start codon, by a cassette consisting of *rev* and *nef* separated by an internal ribosomal entry site (IRES). The *rev* sequence was derived from either the 8-G or 9-G virus.

To create stocks of viral constructs, 293T/17 cells were plated at a density of 3 x 10^6^ cells per 10 cm plate in Iscove’s modified Dulbecco’s media (IMDM) supplemented with 10% bovine calf serum (BCS) and gentamicin. One day after plating, the cells were transfected with 15 μg of plasmid containing one of the replication-competent constructs using the polyethylenamine (PEI) method (48). Supernatant was collected from the transfected cells 48 hours post-transfection, centrifuged briefly to remove cell debris, aliquoted, and stored at −80°C until needed. The concentration of p24 in each transfection stock was determined by ELISA as previously described (49).

Viruses from transfection stocks were expanded by passaging in SupT1 cells. A volume of transfection stock containing 100 ng of p24 was used to infect a culture of 5 x 10^6^ SupT1 cells suspended in serum-free RPMI (PBS) in 50 mL conical tubes. Diethylaminoethyl (DEAE)-dextran was added to each culture to a final concentration of 8 μg/mL and cultures were centrifuged at 25°C at 380 RCF for one hour. After infection, cells were washed with PBS and resuspended in SupT1 growth medium (Roswell Park Memorial Institute 1640 (RPMI) medium supplemented with 10% fetal bovine serum and gentamicin).

Infected SupT1 cultures were serially passaged for up to 33 days. Every 2-3 days, half of the cells and medium were removed from each culture. The collected material was briefly centrifuged to pellet the cells, and the cell-free medium was aliquoted and frozen at −80°C for determination of p24 and for future experiments. After each collection, the culture volume was replenished with sterile SupT1 growth medium. Medium collected on the day of peak p24 was used to determine virus titer by TCID(50) as previously described (50).

### Replication kinetics assay

The replication kinetics of 8-G and 9-G Rev containing viruses was assayed in parallel cultures of SupT1 cells. A total of 3 x 10^5^ SupT1 cells were infected with either 8-G or 9-G Rev containing virus at a multiplicity of infection (MOI) of 0.005. Infections were performed by spinoculation in the presence of 8 μ/mL of DEAE-dextran. After infection, the cell pellet was washed with PBS once and then the cells were suspended in 10 mL SupT1 growth medium, transferred to flasks, and placed in an incubator. On days 0, 1, 3, 5, 8, and 10 after infection, 1 mL of medium with suspended cells was removed from each flask and replaced with 1 mL of sterile SupT1 growth medium. The sample was centrifuged to pellet cells, and the cell-free medium was frozen at −80°C. After all samples were collected, the frozen medium was thawed and the concentration of p24 was determined by ELISA. Data was collected for three independent experiments.

### Competition assay

The contribution of 8-G and 9-G Rev to relative viral fitness was determined by a competition assay. A total of 3 x 10^5^ SupT1 cells were infected with both 8-G and 9-G Rev containing virus at an MOI of 0.005 for each virus stock as above. Infections were performed in triplicate. On days 1, 3, 5, and 8 after infection, 1 mL of medium with suspended cells was removed from each flask and replaced with 1 mL of sterile SupT1 growth medium. The sample was centrifuged to pellet cells, and the cell-free medium was frozen at −80°C. Additionally, a 1:1 ratio of 8G:9G virus mix was used as a day 0 sample to determine initial viral input.

After all time points were collected, viral RNA was extracted from supernatants including the initial virus stock input. Supernatants were centrifuged at 5300 RCF for 10 minutes at 4°C to remove debris, and then supernatant was transferred to a new tube. Viruses were then pelleted by centrifugation at 16000 RCF for 1 h at 4°C and the supernatant was removed and discarded. The virus pellet was resuspended in 50 μL RNAse free 5 mM Tris-HCL, pH 8.0. The suspended viruses were treated with 10 μL proteinase K (20 mg/mL) and incubated at 55°C for 30 minutes. Next, 200 μL 6M guanidinium isothiocyanate and 10 μL glycogen (20 mg/mL) were added to each tube and samples were incubated at 25°C for 5 minutes. RNA was pelleted by adding 270 μL isopropanol to each tube and centrifuging the samples at 2700 RCF for 20 minutes at 25°C. The supernatant was removed, the pellet was washed with 70% ethanol, and then the RNA was resuspended in 40 μL RNAse free Tris-HCL. RT-PCR was performed using the SuperScript III Reverse Transcriptase kit (ThermoFisher) with oligo(dT)20 primer.

The relative proportion of 8-G and 9-G viral RNA was determined by PCR amplification of 2 μL of the prepared cDNA. PCR products from 8-G and 9-G containing viruses were discriminated by size on an agarose gel. PCR was performed using Taq DNA polymerase (Thermo Scientific) per manufacturer recommendations. Each reaction mixture included 2 pmol of a single forward primer located within *env* (5’-CCTAGAAGAATAAGACAGGGC-3’) and 1 pmol each of two reverse primers located within the heterotopic *rev* (8-G specific, 5’-CCCCAGATATTTCAGGCCCTC-3’; 9-G specific, 5’-GTCTCTCAAGCGGTGGTAGCAC-3’). Initial denaturation was performed at 95°C for 3 min; then the reactions were cycled at 94°C for 30 s, 60°C for 30 s, and 75°C for 45 s for 25 cycles; followed by a final extension at 72°C for 10 min. Mixtures of known molar ratios of plasmids containing 8-G and 9-G Rev sequences were used for positive controls.

For each cDNA sample and plasmid standard, 15 uL of PCR product was loaded into each well of a 2% agarose gel containing ethidium bromide and run at 80V for 2 h. Gels were imaged using a ChemiDoc Imager (Biorad) with exposure conditions customized to ensure that no bands were over-saturated. Bands corresponding to 8-G and 9-G Rev targets could be differentiated by size. Gel images were analyzed using Image Lab v 6.1 (Biorad) to quantify the signal from each band.

Lanes corresponding to the known plasmid mixtures were used to construct a standard curve to correlate relative band intensity to molar ratio of 8-G and 9-G Rev target sequence. Data analysis was performed in Excel 2016 (Microsoft). Relative proportion of 8-G and 9-G Rev cDNA was expressed as a percentage of the total signal from both PCR bands in each gel lane.

### Chimeric Rev constructs

Custom rev sequences were designed using either the 8-G or 9-G rev sequence. Functional regions of Rev were defined as in Figure 2. To create chimeras, the named functional region in either the 8-G or 9-G rev RNA sequence was exchanged with the RNA sequence of the corresponding region from 9-G or 8-G, respectively. For example, the “8-G with 9-G ARM+OD” sequence was identical to the 8-G rev sequence except from amino acid positions 35-59 in which region the corresponding sequence from 9-G rev was substituted. At each codon, the nucleotide sequence was taken from either 8-G or 9-G rev. Chimeric sequences and sequences with the specified amino acid substitutions were generated using Geneious Prime v11.0.9 (Biomatters).

### Fluorescence-based functional assays

The fluorescence-based assay of Rev-RRE functional activity, including additional details of the assay constructs, has been previously described (51). The assay system consists of two packageable viral constructs that can be used to co-transduce target lymphoid cells and generate a fluorescent signal in a Rev-RRE dependent fashion.

The Rev-containing vector was derived from a murine stem cell virus construct (pMSCV-IRES-Blue FP), a gift from Dario Vignali (Addgene plasmid #52115). The various *rev* sequences were inserted into this construct upstream of the IRES and fluorescent marker *eBFP2* to permit co-expression from a bicistronic transcript.

The RRE-containing vector was derived from a full-length NL4-3 HIV sequence. The vector was modified in the following ways: native *rev, vpr*, and *env* expression was silenced; and the *gag* myristoylation site was deleted; *gag* was truncated and a cassette including a *cis*-acting hydrolase element from porcine teschovirus-1 (52) and *eGFP* (53) was inserted in-frame; *nef* was deleted and replaced with *mCherry* (54). Additionally, a 350-nt region within *env* containing the RRE was flanked by restriction sites for easy exchange, and assay constructs were created by replacing this region with the 234-nt RRE from the 8-G, 9-G, or NL4-3 sequences.

To create stocks of pseudotyped, packaged assay constructs, 293T/17 cells were plated at a density of 3 x 10^6^ cells per 10 cm plate in IMDM supplemented with 10% BCS and gentamicin. One day after plating, the cell medium was replaced with IMDM supplemented with 5% BCS. The cells were then transfected with 15 ug of the desired Rev- or RRE-containing construct, 2.54 ug of the VSV-G expression plasmid pMD2.G (a gift from Didier Trono, Addgene plasmid # 12259), and 6.42 ug of the appropriate helper construct (pHit-CMV-GagPol (55) for Rev-containing constructs and psPAX2 for RRE-containing constructs) using the PEI method. The plasmid psPAX2 was a gift from Didier Trono (Addgene plasmid # 12260). Medium was collected from the producer culture 48 hours after transfection, aliquoted, and frozen at −80°C until use.

Transduction for vector titering and functional assays was performed in 96 well plates. A total of 2.5 x 10^5^ SupT1 cells suspended in PBS were seeded in each well along with stocks of vectors. Sterile DEAE-dextran was added to each well to a final concentration of 8 mcg/mL. The transduction cultures were then centrifuged at 380 RCF for 1 h at room temperature. After centrifugation, vector-containing medium was removed from the cell pellets, and cells were resuspended in SupT1 growth medium. Seventy-two hours after transduction, cells were again centrifuged briefly to pellet them, and the medium was removed. Cells were resuspended in cold PBS and held on ice in preparation for flow cytometry.

Transductions for determining the titer of vector stocks were performed using serial 1:10 dilutions of the initial stock. Transductions for the functional assays were performed using both Rev- and RRE-containing vectors at a target MOI of 0.18. Transductions for each Rev-RRE pair included in an experimental run were performed in duplicate or triplicate in different wells.

Flow cytometry was performed using an Attune NxT flow cytometer with autosampler attachment (Thermo Fischer Scientific). Data was acquired using the Attune NxT software package using the following channels:

|  |  |  |  |
| --- | --- | --- | --- |
| Laser line | 488 | 405 | 561 |
| Emission filter | 530/30 | 440/50 | 620/15 |
| Fluorescent protein | eGFP | eBFP2 | mCherry |

Color compensation and data analysis was performed after acquisition using FlowJo v10.6.1 (FlowJo, LLC). Gates were constructed to identify single SupT1 cells. Using an untransduced SupT1 population, daughter gates were then constructed to define eBFP2 and mCherry positive populations.

For data analysis, only single cells successfully transduced with both a Rev-containing and RRE-containing vector were considered. For this population, the arithmetic mean fluorescent intensity of eGFP and eBFP2 was determined and the ratio of eGFP/eBFP2 was calculated as the measure of relative Rev-RRE functional activity. This calculation was performed for each well in each experimental run.

### Statistical analysis of functional assays

For each experimental run, the ratio of eGFP/BFP2 was calculated for each of the two or three wells transduced with a particular Rev-RRE pair. The activity measurement of each well corresponding to a Rev-RRE pair was averaged to represent a single estimate of Rev-RRE activity for that pair. Repeated measurements of Rev-RRE pair activity performed in different experimental runs on different days were considered to constitute replicates for the purposes of statistical analysis.

Wells in which less than 500 cells were co-transduced with both the Rev-containing and RRE-containing construct were deemed to be uninformative and not included in activity calculations. Wells in which either the Rev-containing or RRE-containing constructs transduced more than 31% of cells were also excluded from further calculations, as a significant number of cells in these wells would contain two or more integrations of the Rev or RRE construct.

While the relative activity of Rev-RRE pairs in comparison to each other was consistent across experimental runs, the absolute ratio of mean eGFP/eBFP2 signal varied from transduction to transduction. Relative Rev activity on a particular RRE was compared using a linear mixed model by restricted maximum likelihood, with individual experimental run as a random effect and Rev as a fixed effect. Analysis was performed using R v4.1.2 and the lme4 (56) and lmerTest packages (57). The *p* value was adjusted for multiple comparisons using the Hochberg method (58). Activity measurements for any Rev-RRE pair represent at least three replicated experimental runs.

### Structural analysis of 8-G and 9-G Rev

Predicted structures of 8-G and 9-G Rev were calculated using the AlphaFold Collab utility (https://colab.research.google.com/github/deepmind/alphafold/blob/main/notebooks/AlphaFold.ipynb) with default settings (31). The computed structures were compared with a previously published crystal structure of HIV-1 NL4-3 Rev (32) using PyMOL Molecular Graphics System, version 2.5.1 (Schrödinger, LLC).

## Acknowledgements

The authors would like to acknowledge the University of Virginia Flow Cytometry Core (RRID: SCR_017829) for assistance with performance of functional assays and Clay Ford of the University of Virginia Library for assistance with statistical analysis. P. E. H. J. was supported by grant K08AI136671 from the National Institutes of Health. Salary support for G.D. and D.R. was provided by the Myles H. Thaler Research Fund and Professorship at the University of Virginia. Salary support for M.-L. H. was provided by the Charles H. Ross Jr. Professorship at the University of Virginia. Research on HIV Rev in the laboratories of D.R. and M-L. H. was supported by grants CA206275 and AI134208 from the National Institutes of Health.

## Author contributions

P. E. H. J. contributed to conceptualization of the study, performed functional assays, and wrote the draft of the manuscript. G. D. contributed to the conceptualization of the study and performed competition assays. J. H. performed functional assays, E. H. performed PCR for competition assays, and H. O. performed structural analyses. D.R. and M.-L. H contributed to the conceptualization of the studies and experimental design and helped revise the manuscript. All authors reviewed the final manuscript.

## Data availability

The datasets generated during the current study and plasmids used to perform assays are available from the corresponding author on reasonable request.

## Supplemental Material

**Figure S1. Activity of Revs with c-terminus modifications.**

**Figure S2. Activity of Revs with single amino acid substitutions on 8-G and 9-G RREs.**

**Figure S3. Activity of Revs with multiple amino acid substitutions on 8-G and 9-G RREs.**

**Figure S4. Activity of Revs with phosophomimetic substitutions on 8-G and 9-G RREs.**

**Figure S5. Activity of position 19 mutants on 8-G and 9-G RREs.**

**Table S1. List of Rev assay constructs.**

**Table S2. List of RRE assay constructs.**

**Table S3. List of replication-competent constructs.**

## References

1. Rekosh D, Hammarskjold M-L. Intron retention in viruses and cellular genes: Detention, border controls and passports. Wiley Interdisciplinary Reviews: RNA. 2018;9(3):e1470.

2. Frankel AD, Young JA. HIV-1: fifteen proteins and an RNA. Annual Review of Biochemistry. 1998;67(1):1–25.

3. Pollard VW, Malim MH. The HIV-1 rev protein. Annual Reviews in Microbiology. 1998;52(1):491–532.

4. Purcell DF, Martin MA. Alternative splicing of human immunodeficiency virus type 1 mRNA modulates viral protein expression, replication, and infectivity. Journal of virology. 1993;67(11):6365–78.

5. Schwartz S, Felber BK, Benko DM, Fenyo EM, Pavlakis GN. Cloning and functional analysis of multiply spliced mRNA species of human immunodeficiency virus type 1. Journal of virology. 1990;64(6):2519–29.

6. Sadaie MR, Benaissa ZN, Cullen BR, Wong-Staal F. Human immunodeficiency virus type 1 rev protein as a negative trans-regulator. DNA. 1989;8(9):669–74.

7. Hammarskjold ML, Heimer J, Hammarskjold B, Sangwan I, Albert L, Rekosh D. Regulation of human immunodeficiency virus env expression by the rev gene product. J Virol. 1989;63(5):1959–66.

8. Malim MH, Hauber J, Le SY, Maizel JV, Cullen BR. The HIV-1 rev trans-activator acts through a structured target sequence to activate nuclear export of unspliced viral mRNA. Nature. 1989;338(6212):254–7.

9. Daly TJ, Cook KS, Gray GS, Maione TE, Rusche JR. Specific binding of HIV-1 recombinant Rev protein to the Rev-responsive element in vitro. Nature. 1989;342(6251):816–9.

10. Fernandes J, Jayaraman B, Frankel A. The HIV-1 Rev response element: an RNA scaffold that directs the cooperative assembly of a homo-oligomeric ribonucleoprotein complex. RNA biology. 2012;9(1):6–11.

11. Fernandes JD, Booth DS, Frankel AD. A structurally plastic ribonuceloprotein complex mediates post-transcriptional gene regulation in HIV-1. Wiley Interdiscip Rev RNA. 2016.

12. Bai Y, Tambe A, Zhou K, Doudna JA. RNA-guided assembly of Rev-RRE nuclear export complexes. Elife. 2014;3:e03656.

13. Neville M, Stutz F, Lee L, Davis LI, Rosbash M. The importin-beta family member Crm1p bridges the interaction between Rev and the nuclear pore complex during nuclear export. Current Biology. 1997;7(10):767–75.

14. Smyth RP, Davenport MP, Mak J. The origin of genetic diversity in HIV-1. Virus research. 2012;169(2):415–29.

15. Jackson PE, Tebit DM, Rekosh D, Hammarskjold M-L. Rev–RRE Functional Activity Differs Substantially Among Primary HIV-1 Isolates. AIDS research and human retroviruses. 2016.

16. Sloan EA, Kearney MF, Gray LR, Anastos K, Daar ES, Margolick J, et al. Limited nucleotide changes in the Rev response element (RRE) during HIV-1 infection alter overall Rev-RRE activity and Rev multimerization. J Virol. 2013;87(20):11173–86.

17. Sherpa C, Jackson PEH, Gray LR, Anastos K, Le Grice SFJ, Hammarskjold M-L, et al. Evolution of the HIV-1 Rev Response Element during Natural Infection Reveals Nucleotide Changes That Correlate with Altered Structure and Increased Activity over Time. Journal of Virology. 2019;93(11).

18. Sherpa C, Rausch JW, Le Grice SFJ, Hammarskjold M-L, Rekosh D. The HIV-1 Rev response element (RRE) adopts alternative conformations that promote different rates of virus replication. Nucleic acids research. 2015;43(9):4676–86.

19. Jain C, Belasco JG. Structural model for the cooperative assembly of HIV-1 Rev multimers on the RRE as deduced from analysis of assembly-defective mutants. Mol Cell. 2001;7(3):603–14.

20. Venkatesh L, Mohammed S, Chinnadurai G. Functional domains of the HIV-1 rev gene required for trans-regulation and subcellular localization. Virology. 1990;176(1):39–47.

21. Heaphy S, Dingwall C, Ernberg I, Gait MJ, Green SM, Karn J, et al. HIV-1 regulator of virion expression (Rev) protein binds to an RNA stem-loop structure located within the Rev response element region. Cell. 1990;60(4):685–93.

22. Kjems J, Frankel AD, Sharp PA. Specific regulation of mRNA splicing in vitro by a peptide from HIV-1 Rev. Cell. 1991;67(1):169–78.

23. Battiste JL, Mao H, Rao NS, Tan R, Muhandiram DR, Kay LE, et al. Alpha helix-RNA major groove recognition in an HIV-1 rev peptide-RRE RNA complex. Science. 1996;273(5281):1547–51.

24. Fischer U, Huber J, Boelens WC, Mattaj IW, Luhrmann R. The HIV-1 Rev activation domain is a nuclear export signal that accesses an export pathway used by specific cellular RNAs. Cell. 1995;82(3):475–83.

25. Jayaraman B, Fernandes J, Yang S, Smith C, Frankel AD. Highly Mutable Linker Regions Regulate HIV-1 Rev Function and Stability. bioRxiv. 2018:424259.

26. Jackson PE, Huang J, Sharma M, Rasmussen SK, Hammarskjold M-L, Rekosh D. A novel retroviral vector system to analyze expression from mRNA with retained introns using fluorescent proteins and flow cytometry. Scientific reports. 2019;9(1):1–14.

27. Meggio F, D’Agostino DM, Ciminale V, Chieco-Bianchi L, Pinna LA. Phosphorylation of HIV-1 Rev protein: implication of protein kinase CK2 and pro-directed kinases. Biochemical and biophysical research communications. 1996;226(2):547–54.

28. Marin O, Sarno S, Boschetti M, Pagano MA, Meggio F, Ciminale V, et al. Unique features of HIV-1 Rev protein phosphorylation by protein kinase CK2 (‘casein kinase-2’). FEBS Letters. 2000;481(1):63–7.

29. Fouts DE, True HL, Cengel KA, Celander DW. Site-specific phosphorylation of the human immunodeficiency virus type-1 Rev protein accelerates formation of an efficient RNA-binding conformation. Biochemistry. 1997;36(43):13256–62.

30. Chen Z, Cole PA. Synthetic approaches to protein phosphorylation. Current opinion in chemical biology. 2015;28:115–22.

31. Jumper J, Evans R, Pritzel A, Green T, Figurnov M, Ronneberger O, et al. Highly accurate protein structure prediction with AlphaFold. Nature. 2021;596(7873):583–9.

32. Jayaraman B, Crosby DC, Homer C, Ribeiro I, Mavor D, Frankel AD. RNA-directed remodeling of the HIV-1 protein Rev orchestrates assembly of the Rev–Rev response element complex. Elife. 2015;3:e04120.

33. Bobbitt KR, Addo MM, Altfeld M, Filzen T, Onafuwa AA, Walker BD, et al. Rev activity determines sensitivity of HIV-1-infected primary T cells to CTL killing. Immunity. 2003;18(2):289–99.

34. Phuphuakrat A, Auewarakul P. Functional variability of Rev response element in HIV-1 primary isolates. Virus genes. 2005;30(1):23–9.

35. Phuphuakrat A, Paris RM, Nittayaphan S, Louisirirotchanakul S, Auewarakul P. Functional variation of HIV-1 Rev response element in a longitudinally studied cohort. Journal of medical virology. 2005;75(3):367–73.

36. Oelrichs R, Tsykin A, Rhodes D, Solomon A, Ellett A, McPhee D, et al. Genomic sequence of HIV type 1 from four members of the Sydney Blood Bank Cohort of long-term nonprogressors. AIDS research and human retroviruses. 1998;14(9):811.

37. Papathanasopoulos MA, Patience T, Meyers TM, McCutchan FE, Morris L. Full-length genome characterization of HIV type 1 subtype C isolates from two slow-progressing perinatally infected siblings in South Africa. AIDS Research & Human Retroviruses. 2003;19(11):1033–7.

38. Edgcomb SP, Aschrafi A, Kompfner E, Williamson JR, Gerace L, Hennig M. Protein structure and oligomerization are important for the formation of export-competent HIV-1 Rev-RRE complexes. Protein Science. 2008;17(3):420–30.

39. Svicher V, Alteri C, D’sArrigo R, Lagana A, Trignetti M, Lo Caputo S, et al. Treatment with the fusion inhibitor enfuvirtide influences the appearance of mutations in the human immunodeficiency virus type 1 regulatory protein rev. Antimicrobial agents and chemotherapy. 2009;53(7):2816–23.

40. Hua J, Caffrey JJ, Cullen BR. Functional consequences of natural sequence variation in the activation domain of HIV-1 Rev. Virology. 1996;222(2):423–9.

41. Iversen AK, Shpaer EG, Rodrigo AG, Hirsch MS, Walker BD, Sheppard HW, et al. Persistence of attenuated rev genes in a human immunodeficiency virus type 1-infected asymptomatic individual. J Virol. 1995;69(9):5743–53.

42. Churchill MJ, Chiavaroli L, Wesselingh SL, Gorry PR. Persistence of attenuated HIV-1 rev alleles in an epidemiologically linked cohort of long-term survivors infected with nef-deleted virus. Retrovirology. 2007;4:43.

43. Fernandes JD, Faust TB, Strauli NB, Smith C, Crosby DC, Nakamura RL, et al. Functional Segregation of Overlapping Genes in HIV. Cell. 2016;167(7):1762–73.e12.

44. Yamaguchi J, Ndembi N, Ngansop C, Mbanya D, Kaptue L, Gürtler LG, et al. HIV type 1 group M subtype G in Cameroon: five genome sequences. AIDS research and human retroviruses. 2009;25(4):469–73.

45. Sanchez AM, DeMarco CT, Hora B, Keinonen S, Chen Y, Brinkley C, et al. Development of a contemporary globally diverse HIV viral panel by the EQAPOL program. Journal of immunological methods. 2014;409:117–30.

46. Salminen MO, Koch C, Sanders-Buell E, Ehrenberg PK, Michael NL, Carr JK, et al. Recovery of virtually full-length HIV-1 provirus of diverse subtypes from primary virus cultures using the polymerase chain reaction. Virology. 1995;213(1):80–6.

47. Adachi A, Gendelman HE, Koenig S, Folks T, Willey R, Rabson A, et al. Production of acquired immunodeficiency syndrome-associated retrovirus in human and nonhuman cells transfected with an infectious molecular clone. J Virol. 1986;59(2):284–91.

48. Boussif O, Lezoualc’h F, Zanta MA, Mergny MD, Scherman D, Demeneix B, et al. A versatile vector for gene and oligonucleotide transfer into cells in culture and in vivo: polyethylenimine. Proceedings of the National Academy of Sciences of the United States of America. 1995;92(16):7297–301.

49. Wehrly K, Chesebro B. p24 antigen capture assay for quantification of human immunodeficiency virus using readily available inexpensive reagents. Methods. 1997;12(4):288–93.

50. Gao Y, Nankya I, Abraha A, Troyer RM, Nelson KN, Rubio A, et al. Calculating HIV-1 Infectious Titre Using a Virtual TCID 50 Method. HIV protocols: Springer; 2008. p. 27–35.

51. Jackson PEHH, Jing; Sharma, Monika; Rasmussen, Sara K.; Hammarskjold, Marie-Louise; Rekosh, David. A novel fluorescence-based assay to measure the activity of RNA elements that promote the export of mRNA with retained introns, using packageable HIV-based vectors. BioRXiv. 2019.

52. Kim JH, Lee SR, Li LH, Park HJ, Park JH, Lee KY, et al. High cleavage efficiency of a 2A peptide derived from porcine teschovirus-1 in human cell lines, zebrafish and mice. PLoS One. 2011;6(4):e18556.

53. Zhang G, Gurtu V, Kain SR. An enhanced green fluorescent protein allows sensitive detection of gene transfer in mammalian cells. Biochem Biophys Res Commun. 1996;227(3):707–11.

54. Shaner NC, Campbell RE, Steinbach PA, Giepmans BN, Palmer AE, Tsien RY. Improved monomeric red, orange and yellow fluorescent proteins derived from Discosoma sp. red fluorescent protein. Nat Biotechnol. 2004;22(12):1567–72.

55. Soneoka Y, Cannon PM, Ramsdale EE, Griffiths JC, Romano G, Kingsman SM, et al. A transient three-plasmid expression system for the production of high titer retroviral vectors. Nucleic acids research. 1995;23(4):628–33.

56. Bates D, Maechler M, Bolker B, Walker S. Fitting linear mixed-effects models using lme4. Journal of Statistical Software 67:1–48. 2015.

57. Kuznetsova A, Brockhoff PB, Christensen RH. lmerTest package: tests in linear mixed effects models. Journal of statistical software. 2017;82:1–26.

58. Hochberg Y. A sharper Bonferroni procedure for multiple tests of significance. Biometrika. 1988;75(4):800–2.

