## Supplemental material for "HIV-1 Rev-RRE Functional Activity in Primary Isolates is Highly Dependent on Minimal Context-Dependent Changes in Rev"

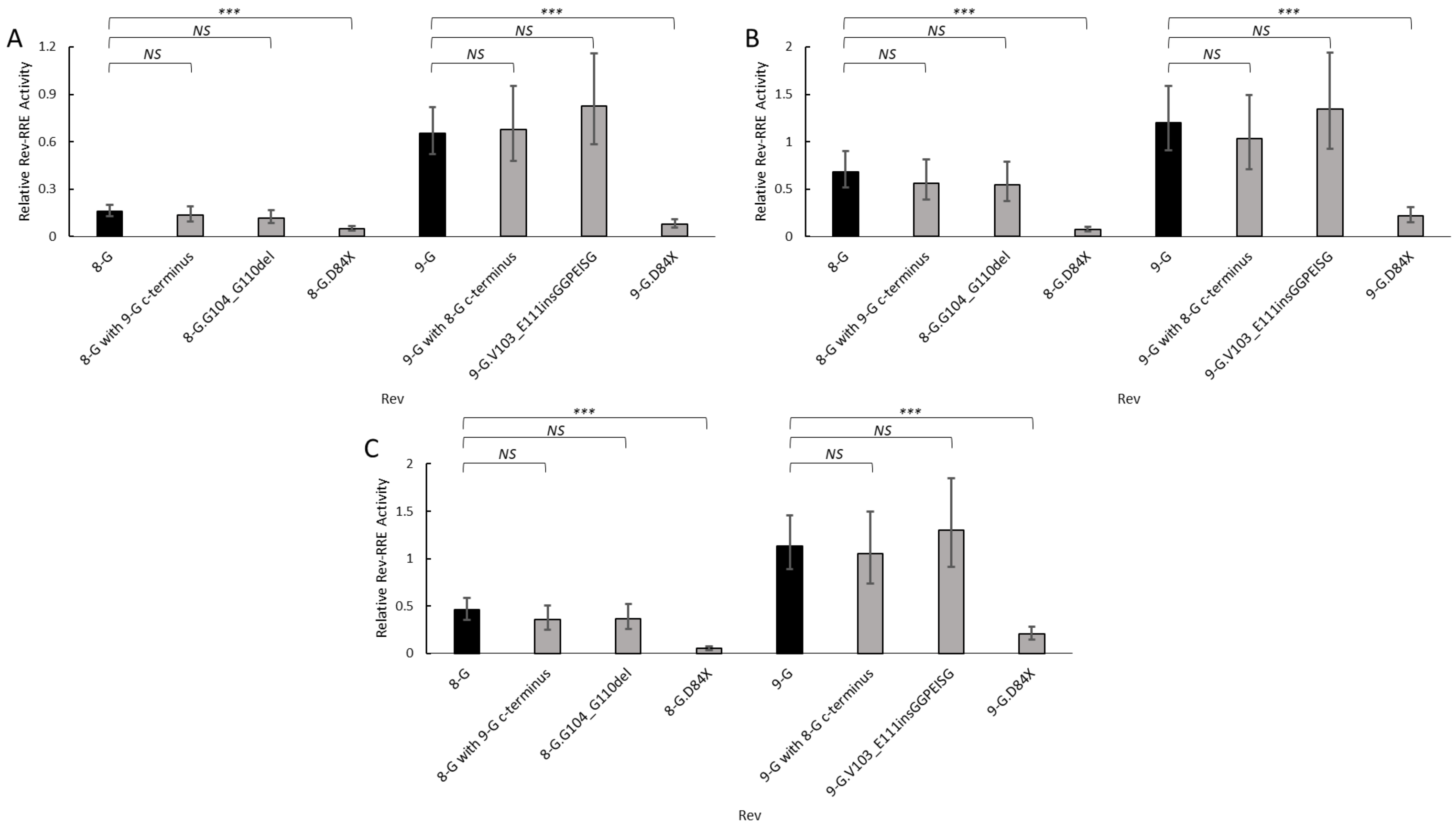


**Figure S1. Activity of Revs with c-terminus modifications.**

8-G and 9-G Revs were created with modifications to the c-terminal region and their functional activity was determined using the fluorescence-based assay system. In each plot, the native 8-G and 9-G Rev sequences are included in dark bars for reference and modified Revs are in light bars. The Revs were tested with the NL4-3 RRE (A), 8-G RRE (B), and 9-G RRE (C). *N* ≥ 3 for all data points, error bars represent 95% CI. *NS* – not significant, *** $p < 0.001$.

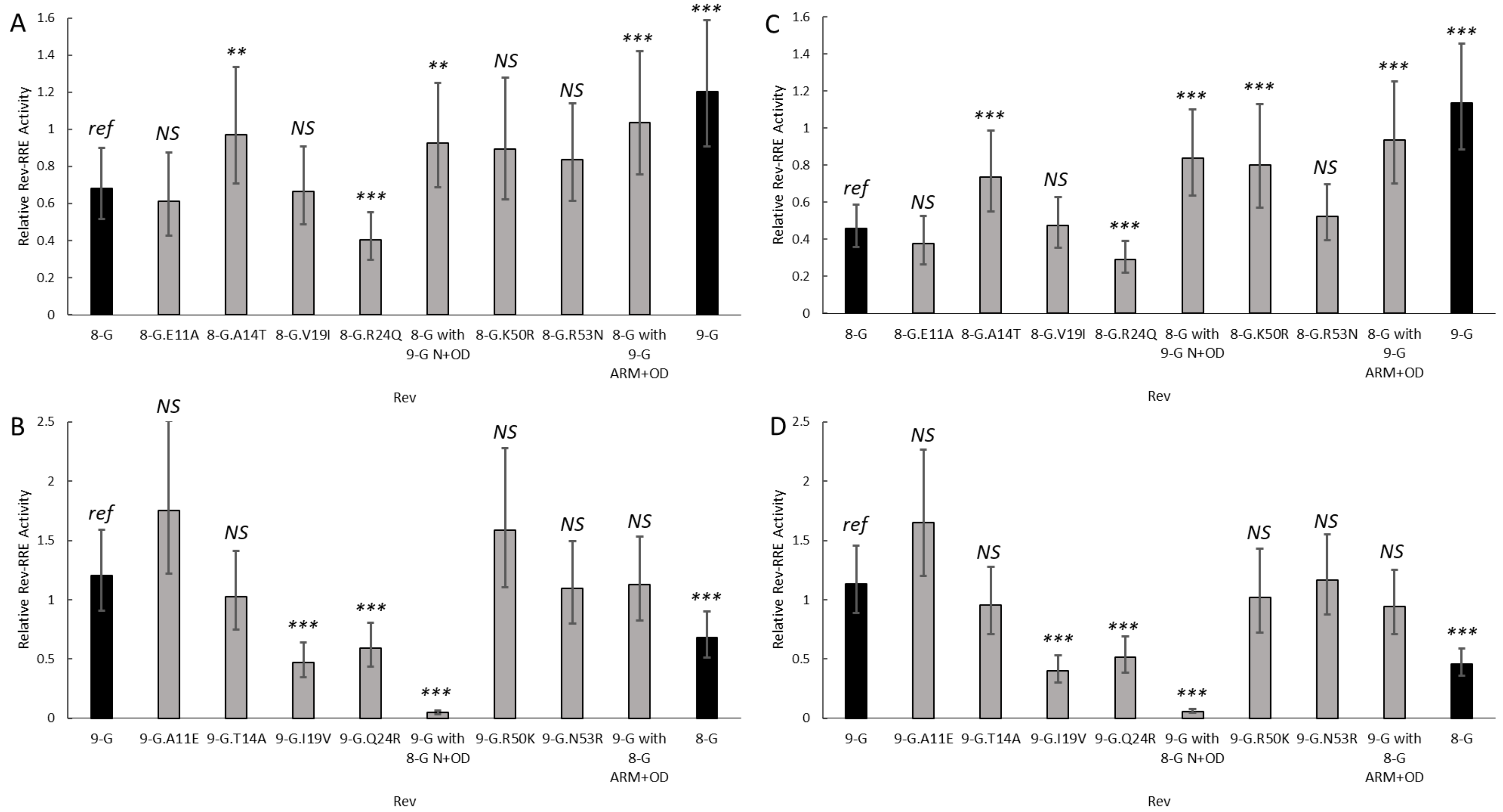


**Figure S2. Activity of Revs with single amino acid substitutions on 8-G and 9-G RREs.**

Revs with single amino acid substitutions within the N-terminal, OD, or ARM regions were created and their functional activity determined using the fluorescence-based assay system. A. 8-G derived Revs tested on the 8-G RRE. B. 9-G derived Revs tested on the 8-G RRE. C. 8-G derived Revs tested on the 9-G RRE. D. 9-G derived Revs tested on the 9-G RRE. In each plot, the native 8-G and 9-G Rev sequences are included in dark bars for reference and modified Revs are in light bars. The statistical comparison is performed in reference to the left-most Rev in each plot. $N \geq 3$ for all data points, error bars represent 95% CI. *Ref* – reference sequence for the plot, *NS* – not significant, *** $p < 0.001$, ** $p < 0.01$.

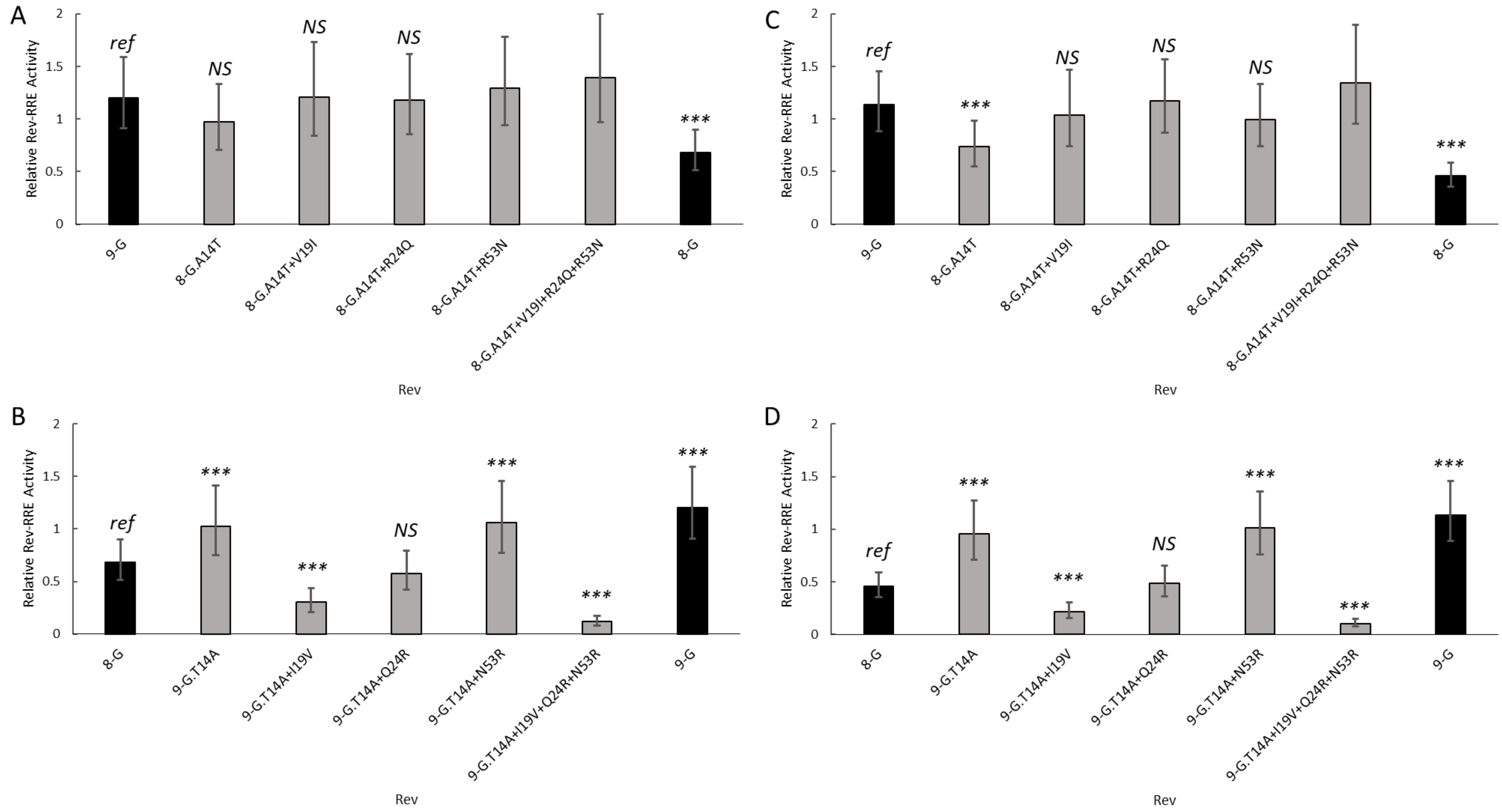


**Figure S3. Activity of Revs with multiple amino acid substitutions on 8-G and 9-G RREs.**

Revs with multiple amino acid substitutions were created and their functional activity determined using the fluorescence-based assay system. A. 8-G derived Revs tested on the 8-G RRE. B. 9-G derived Revs tested on the 8-G RRE. C. 8-G derived Revs tested on the 9-G RRE. D. 9-G derived Revs tested on the 9-G RRE. In each plot, the native 8-G and 9-G Rev sequences are included in dark bars for reference and modified Revs are in light bars. The statistical comparison is performed in reference to the left-most Rev in each plot. $N \geq 3$ for all data points, error bars represent 95% CI. *Ref* – reference sequence for the plot, *NS* – not significant, *** $p < 0.001$.

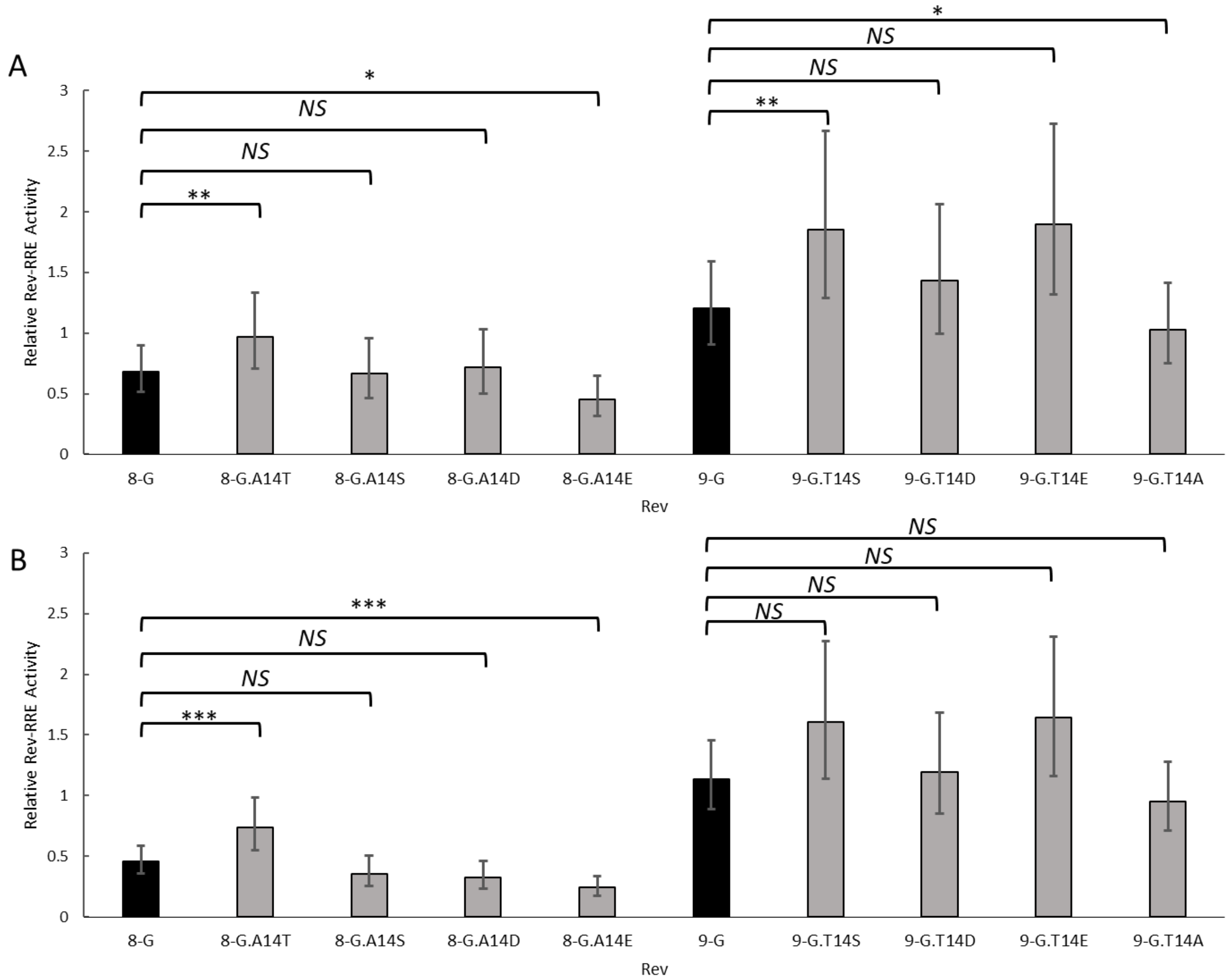


**Figure S4. Activity of Revs with phosophomimetic substitutions on 8-G and 9-G RREs.**

Revs with phosphomimetic amino acid substitutions at position 14 were created and their functional activity determined on the 8-G RRE (A) and 9-G RRE (B) using the fluorescence-based assay system. The native 8-G and 9-G Rev sequences are included in dark bars for reference and modified Revs are in light bars. The statistical comparison is performed is performed between the Revs indicated by the horizontal bars. $N \geq 3$ for all data points, error bars represent 95% CI. *NS* – not significant, *** $p < 0.001$, ** $p < 0.01$, * $p < 0.05$.

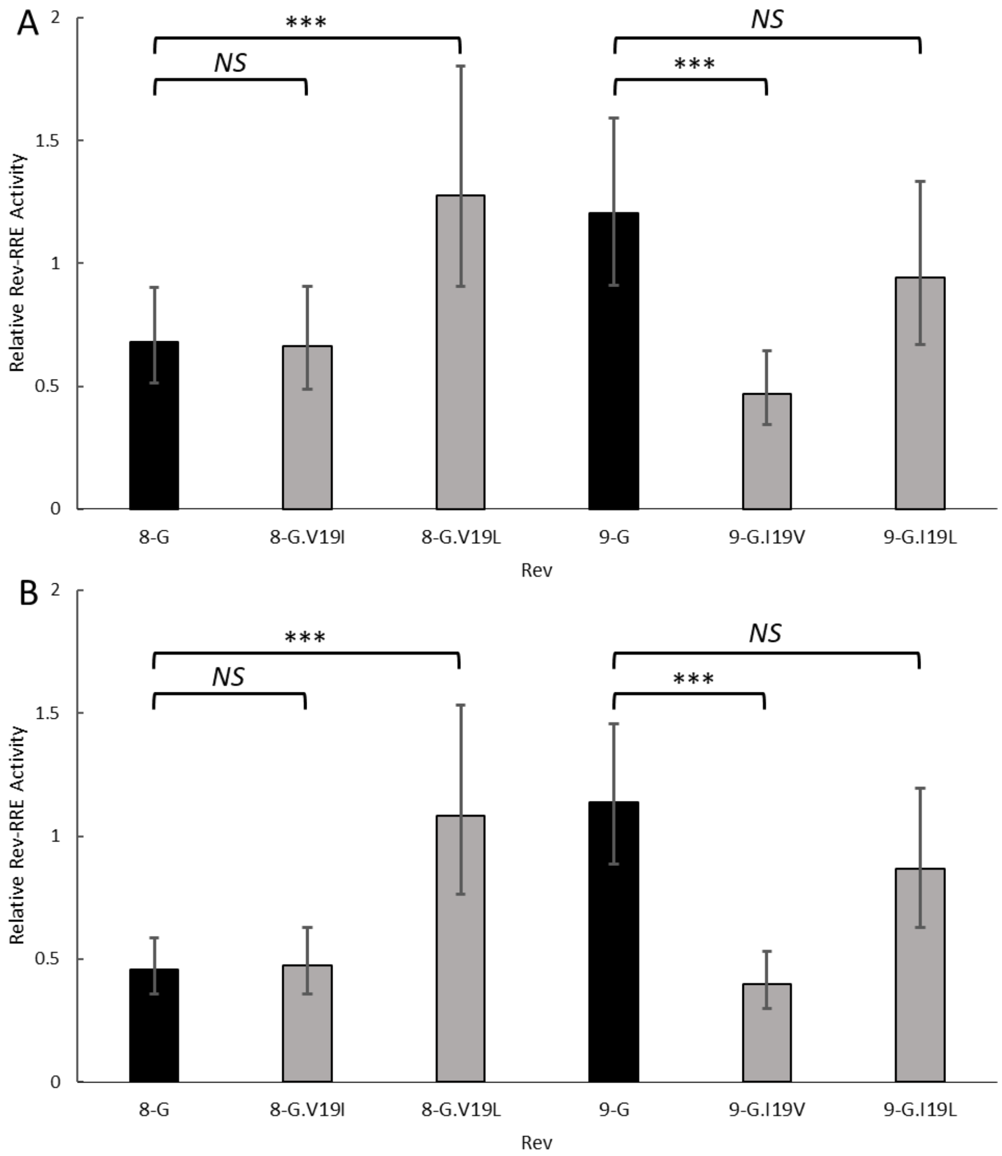


**Figure S5. Activity of position 19 mutants on 8-G and 9-G RREs.**

Revs with amino acid substitutions at position 10 were created and their functional activity determined on the 8-G RRE (A) and 9-G RRE (B) using the fluorescence-based assay system. The native 8-G and 9-G Rev sequences are included in dark bars for reference and modified Revs are in light bars. The statistical comparison is performed is performed between the Revs indicated by the horizontal bars. $N \geq 3$ for all data points, error bars represent 95% CI. *NS* – not significant, *** $p < 0.001$.

**Table S1. List of Rev assay constructs.**

| Rev construct | Plasmid number |
|---|---|
| NL4-3 | 5864 |
| 9-G | 5866 |
| 8-G | 5868 |
| 8-G with 9-G Link | 5912 |
| 8-G with 9-G Turn | 5914 |
| 8-G with 9-G c-terminus | 5916 |
| 8-G.R24Q | 5918 |
| 9-G with 8-G Link | 5920 |
| 9-G with 8-G Turn | 5922 |
| 9-G with 8-G c-terminus | 5924 |
| 9-G.Q24R | 5926 |
| 8-G.G104_G110del | 5928 |
| 9-G.V103_E111insGGPEISG | 5930 |
| 8-G with 9-G ARM+OD | 5982 |
| 8-G with 9-G N+OD | 5984 |
| 8-G with 9-G NES | 5986 |
| 9-G with 8-G ARM+OD | 5988 |
| 9-G with 8-G N+OD | 5990 |
| 9-G with 8-G NES | 5992 |
| NL4-3.Q24R | 5994 |
| 8-G.A14T | 6080 |
| 8-G.D84X | 6081 |
| 8-G.E11A | 6082 |
| 8-G.V19I+R24Q | 6083 |
| 8-G.V19I | 6084 |
| 9-G.A11E | 6085 |
| 9-G.D84X | 6086 |
| 9-G.I19V+Q24R | 6087 |
| 9-G.I19V | 6088 |
| 9-G.T14A | 6089 |
| 8-G.A14T+R24Q | 6105 |
| 8-G.A14T+R53N | 6106 |
| 8-G.A14T+V19I | 6107 |

| | |
|---|---|
| 8-G.A14T+V19I+R24Q+R53N | 6108 |
| 8-G.K50R | 6109 |
| 8-G.R53N | 6110 |
| 9-G.N53R | 6111 |
| 9-G.R50K | 6112 |
| 9-G.T14A+I19V | 6113 |
| 9-G.T14A+I19V+Q24R+N53R | 6114 |
| 9-G.T14A+N53R | 6115 |
| 9-G.T14A+Q24R | 6116 |
| 9-G.I19L+N53R | 6666 |
| 9-G.I19L | 6667 |
| 8-G.V19L+R53N | 6668 |
| 8-G.V19L | 6669 |

**Table S2. List of RRE assay constructs.**

| **RRE construct** | **Plasmid number** |
|---|---|
| NL4-3 | 5936 |
| 8-G | 5938 |
| 9-G | 5940 |

**Table S3. List of replication-competent constructs.**

| **Construct** | **Plasmid number** |
|---|---|
| pNL4-3(Rev-)(8-G Rev-IRES-Nef) | 5830 |
| pNL4-3(Rev-)(9-G Rev-IRES-Nef) | 5831 |
